# Transcription and DNA methylation signatures of paternal behavior in hippocampal dentate gyrus of prairie voles

**DOI:** 10.1101/2022.09.09.507382

**Authors:** Nicholas J. Waddell, Yan Liu, Javed M. Chitaman, Graham J. Kaplan, Zuoxin Wang, Jian Feng

## Abstract

In socially monogamous prairie voles (*Microtus ochrogaster*), parental behaviors not only occur in mothers and fathers, but also exist in some virgin males. In contrast, the other virgin males display aggressive behaviors towards conspecific pups. However, little is known about the molecular underpinnings of this behavioral dichotomy, such as gene expression changes and their regulatory mechanisms. To address this, we profiled the transcriptome and DNA methylome of hippocampal dentate gyrus of four prairie vole groups, namely attacker virgin males, parental virgin males, fathers, and mothers. While we found a concordant gene expression pattern between parental virgin males and fathers, the attacker virgin males have a more deviated transcriptome. Moreover, numerous DNA methylation changes were found in pair-wise comparisons among the four groups. We found some of these DNA methylation changes correlate with transcription differences, particularly at genes with extreme expression changes. Furthermore, the gene expression changes and methylome alterations are selectively enriched in certain biological pathways, such as Wnt signaling, which suggest a canonical transcription regulatory role of DNA methylation in paternal behavior. Therefore, our study presents an integrated view of prairie vole dentate gyrus transcriptome and epigenome that provides a DNA epigenetic based molecular insight of paternal behavior.

## INTRODUCTION

In mothers, gene expression changes in the brain underlie the physiological and behavioral adaptations of pup-rearing [1]. In addition, accumulating evidence suggests that neural gene expression changes are associated with sexual experience in fathers of biparental species, where both parents participate in pup-rearing, and paternal care contributes to pup development [2]. However, the regulatory mechanism of gene expression underlying these processes remains elusive.

The prairie vole, *Microtus ochrogaster*, has become a valuable organism to model social bonding, where vole individuals mate exclusively, share nests, and exhibit biparental care of newborn pups [3–5]. Both mother and father voles nearly equally participate in parental behaviors (“Mother” and “Father” thereafter, respectively), such as pup grooming, huddling, retrieving, and nest building [6]. Though these parental behaviors normally present after a litter is born, they may spontaneously occur in some sexually naïve males when exposed to conspecific pups (“Parental”) [7, 8]. While about 60% virgin males are parental, the other virgin males display aggression towards conspecific pups (“Attacker”) [9–11]. Therefore, to study the behavioral dichotomy in male prairie voles may contribute to our understanding of paternal care.

Environmental stimuli play a large role in parental behavior manifestation and modulation during post-partum periods, which also affect gene expression through epigenetic modulations [12, 13]. For example, it was demonstrated that parental behaviors in response to pup exposure can be altered by using compounds affecting epigenetic states, such as histone deacetylase inhibitors [14]. Furthermore, differential DNA methylation has been observed in the hippocampus of rodent offspring upon altered maternal care [15]. Although DNA methylation, a major epigenetic mechanism, has been implicated in various basic brain functions and diseases [16–18], the potential role of DNA methylation in paternal behaviors remains largely unknown.

In prairie voles, dentate gyrus (DG) of hippocampus expresses receptors for oxytocin [19], a key molecule in pair bonding and parental behaviors. It was found that oxytocin receptors are subjected to DNA methylation mediated gene expression regulation [20], whose density in DG is associated with mating tactics and reproductive success in male voles [21]. Furthermore, mating and social interaction, which lead to pair bond formation, have been found to modulate neural precursor cell proliferation and differentiation in the DG of parental voles [22, 23]. While pup exposure elicited cell proliferation [24], fatherhood decreased cell survival in the DG [25]. In addition, exposure to psychostimulant drugs, such as amphetamine, not only diminished pair bonding [26, 27], but also impaired social recognition and decreased neuronal and neurochemical activation in the DG [28]. Although these sporadic evidence accumulatively indicate a role of DNA methylation in DG’s function in prairie vole social behaviors, it remains unknown how gene expression and DNA methylation changes occur at the genomic scale and their potential interplay in parental behaviors. To address this question, we examined the prairie vole DG transcriptome and DNA methylome aiming to explore molecular insights of parental behaviors, particularly the paternal behavioral dichotomy in virgin males.

## METHODS

### Animal Subjects

Subjects were sexually-naïve male and female prairie voles, *Microtus ochrogaster*, from a laboratory breeding colony. Subjects were weaned at 21 days of age and housed in same-sex sibling pairs in plastic cages (12 W x 28 L x 16 H cm) containing cedar chip bedding with water and food provided ad libitum. All cages were maintained under a 14:10 light: dark cycle, and the temperature was kept at 20°C. Adult subjects (at 90-120 days of age) were randomly assigned into experimental groups in which males and females were paired or virgin males were continuously housed in same-sex sibling pairs. Females gave birth following 21-23 days of pairing with a male, and the mother and father voles were continuously housed with their offspring. At three days postpartum, mothers and fathers (i.e. “Mother” and “Father” groups, respectively) were tested for their parental behaviors towards a conspecific pup. Age-matched virgin males were also tested for their spontaneous parental behaviors towards a conspecific pup. As virgin males either display parental behaviors towards pups or attack pups [10, 29, 30]), these males were classified into “Parental” and “Attacker” groups, respectively. All animal experimental procedures were approved by the Florida State University Institutional Animal Care and Use Committee and were in accordance with the U.S. National Institutes of Health Guide for the Care and Use of Laboratory Animals [31].

### Parental Behavior Test

The parental behavior test was conducted as previously described [10, 29, 30]. Briefly, all subjects were tested in a plexiglas cage (20 W x 45 L x 25 H cm) with a thin layer of cedar chip bedding and ad lib food and water, as described for the housing cages. The subject was placed in the testing cage and allowed for a 15-min habituation. Afterwards, an unfamiliar stimulus pup (at 3-day age) was introduced into the testing cage at the opposite corner from the subject, and the subject’s behaviors were digitally recorded for 60 minutes. If the virgin male subject showed aggression towards the pup, the experimenter immediately stopped the behavioral testing by removing the pup, and the subject was classified as “Attacker”.

All behavioral videos were scored by a trained experimenter blind to the treatment groups using JWatcher software program v1.0 [32]. The duration and frequency of the subject’s interactions with the stimulus pup within the first 10-min were quantified. The scored behaviors included both parental behaviors (pup huddling, pup carrying, licking and grooming, and nest building) and non-parental behaviors (auto-grooming, locomotion, olfaction, and resting) [8, 29, 33]. For virgin male attackers, only the latency of the first attack was scored.

### Behavior Data Analysis

Group differences in all behavioral measurements were analyzed using a one-way ANOVA. Post-hoc analyses were conducted using Tukey’s HSD tests (p < 0.05). Plots were generated from ggplot2 v3.3.3 [34].

### Brain Tissue Collection

After the parental behavioral test, subjects were decapitated. Brains were extracted and immediately frozen on dry ice. Brains were sliced into 200 µm sections on a cryostat and thaw mounted on slides. Thereafter, 1 mm-diameter punches from 4 consecutive sections were taken bilaterally from the DG of the hippocampus [35]. Tissue punches were stored at −80°C until further processing.

### Next Generation Sequencing Library Preparation

DNA and RNA were isolated from the same tissue using the Qiagen AllPrep DNA/RNA Micro Kit (Qiagen, #80284) according to the manufacturer’s protocol and then quantified by Qubit Fluorometer (Thermo Fisher Scientific). Each sequencing library was prepared from DNA or RNA isolated from a single mouse brain. Total twenty-three RNAseq libraries (6 Attacker, 6 Parental, 5 Father, 6 Mother) and twenty-three RRBS libraries (6 Attacker, 6 Parental, 5 Father, 6 Mother) were included in the study.

150 ng of total RNA was applied to the NEBNext rRNA Depletion Kit (New England Biolabs, #E6310L) according to the manufacturer’s protocol. Ribo-depleted RNA sequencing (RNA-seq) libraries were constructed using the NEBNext Ultra II Directional RNA Library Prep Kit for Illumina (New England Biolabs, #E7765S). RNA samples were fragmented, and cDNA was synthesized. The ends of cDNA fragments were ligated with universal Illumina adapters. RNAseq libraries were individually indexed with NEBNext Multiplex Oligos for Illumina (New England Biolabs, #E7335S) and amplified for 11 cycles of PCR amplification. All clean-up steps were accomplished by using the supplied purification beads in the NEBNext Ultra II Directional RNA Library Prep Kit. RNAseq libraries were then sequenced 50-bp paired-ended on an Illumina NovaSeq 6000 Sequencer.

Reduced representation bisulfite sequencing (RRBS) [36] libraries were prepared using a Premium RRBS Kit (Diagenode, #C02030032) according to the manufacturer’s protocol. Briefly, for each library preparation, 100 ng genomic DNA was digested with MspI, end-repaired, ligated to adapters, size selected using AMPure XP beads (Beckman Coulter, #A63881), and treated with sodium bisulfite. Spike-in control DNA was included for evaluation of bisulfite conversion efficiency. Libraries were purified after 14 cycles of PCR amplification. Agilent Bioanalyzer and KAPA Library Quantification Kit were applied to assess library quality and quantity. RRBS libraries were sequenced 100-bp single-ended on an Illumina NovaSeq 6000 Sequencer with a 5% PhiX spike-in control.

### Sequencing Data Pre-processing

Raw sequencing reads were first evaluated for quality using FastQC v0.11.9 [37]. To address a positional sequencing error within the RNAseq library, where an entire tile within the sequencing chip had extremely low quality, FilterByTile [38] was applied without incurring biases on the rest of the dataset. Afterwards, sequencing reads were trimmed of adapters and to a minimum of 20 quality score and 20 read length using TrimGalore v0.6.4 [39]. RRBS sequencing reads were trimmed with TrimGalore’s RRBS mode which eliminates synthetic cytosine signals from the ends of reads that was incorporated during the end-repair process. Reads were checked again for quality after trimming, before further analysis.

### RNAseq Analysis

#### Alignment, Assignment, and Differential Expression

Alignment of RNAseq reads was performed using the splice-junction aware alignment software STAR v2.5.4b [40] with the MicOch1.0 [41] annotation set as the reference genome. Aligned reads were assigned and counted to gene-level features using FeatureCounts v2.0.0 [42].

Gene-level counts from RNAseq reads were imported to R and analyzed using EdgeR v3.28.1 [43]. Genes with low counts and those missing counts from at least half of the samples per group were removed from the analysis, resulting in about 70% of annotated genes to be considered in downstream analyses. Dispersion, biological coefficients of variation (BCV) and normalization factors for the dataset were subsequently estimated. RNAseq samples were evaluated in two-dimensional space using multi-dimensional scaling to determine whether certain principal components are driving the variation among and between groups. Then, the RNAseq design matrix and generalized linear models were created contrasting the four groups of voles (“Attacker”, “Parental”, “Father”, and “Mother”) in the experiment. The “Mother” group was only compared to the “Father” group, as both have experienced pair-bonding, while the virgin male vole groups (i.e., “Attacker” and “Parental”) have not, to reduce confounding variations in the analysis. Hypothesis testing was performed through the likelihood ratio test and any genes with a log_2_ fold change less than −0.5 or greater than 0.5, and a p-value <0.05 were claimed significantly differentially expressed. These results were represented using volcano plots generated using ggplots2 v3.3.3 [34].

#### Gene Ontology Enrichment Analysis

Differentially expressed prairie vole genes were annotated to the mouse orthologous gene IDs using Ensembl’s biomart [41] annotation database and SQL manipulations, before they were applied for Kyoto Encylcopedia of Gene and Genomes (KEGG) [44] pathway analysis, which overcomes KEGG’s lack of annotation for prairie voles. The analysis was performed using a web-based tool, WebGestalt [45], a hypergeometric overlaps test. Significant pathways were considered by an FDR value < 0.05. For biological process gene ontology pathway enrichment testing, prairie vole gene names were passed to gProfiler [46] for gene ontology over-representation analysis using Ensembl’s [41] prairie vole annotation. Enrichment ratio is defined by the following formula:

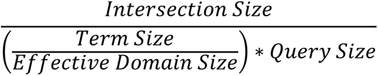

#### Rank-Rank Hypergeometric Overlaps

Rank-rank hypergeometric overlaps (RRHO) analysis identifies overlapping transcriptome expression profiles without pre-set thresholds, and determines the degree and the direction of overlapping genes [47]. An improved version of RRHO is applied to allow discordant signatures to be assessed as robustly as concordant signatures. With this, visualization of each quadrant is separated, where the lengths of each side representing the relative length of each input gene list [48, 49]. Each expression list was ranked by multiplying the −log_10_(p-value) and the sign of the log_2_ fold expression change. RRHO difference maps were generated by representing the −log_10_, Benjamini and Yekutieli adjusted p-value from the hypergeometric test [50].

#### Clustering Analysis

*Differential Expression Clustering*: All genes that had a significant expression change between experimental groups (i.e., “Attacker” vs “Parental”, “Attacker” vs “Father”, “Parental” vs “Father”, and “Father” vs “Mother”; Table S2) were collected, and normalized gene counts were formatted into a table in R. Z-scores were calculated for each gene and passed to the pheatmap v1.0.12 [51] package for hierarchical clustering using k-means clustering based on Euclidean distance. All eight clusters were named in roman numbers I to VIII.

*Gene Ontology Clustering:* Genes in cluster VI of the “Differential Expression Clustering” analysis were taken for evaluation of over-represented ontologies. To simplify the interpretation, the gene ontology pathways were further divided into 10 clusters (named in Arabic numbers 1 to 10), and a network was constructed using Gene Ontology Markov Clustering (GOMCL) [52]. The resulting network and annotation table were passed to Cytoscape v3.9.1 [53] for network visualization.

### RRBS Analysis

#### Alignment, Methylation Calling, Differential Methylation Analysis

Bismark v22.3 [54] genome preparation tool was used to create appropriate reference genomes for bisulfite sequencing alignment. Subsequently, quality-checked RRBS sequencing reads were aligned using the bowtie2 based methylation alignment algorithm in the Bismark suite with increased seed extension effort (parameters: -N 1, -L 20, -D 20). To extract methylation status from the alignment data, a methylation extractor tool of Bismark is applied to create a sample-wise list of CpG positions with the number of reads at that location with methylated calls and unmethylated calls. The resulting files were formatted and imported to R for differential methylation analysis. Differential methylation was calculated using a similar design structure to the differential expression analysis, which was processed using DSS general v2.34.0 [55] that implements a bayesian hierarchical model for dispersion estimation of the beta binomial distribution. Differentially methylated CpG sites were those from the Wald testing procedure with p-value thresholding < 0.05 and an absolute methylation difference of 15%. The data was represented by a volcano plot created using ggplot2 [34].

#### Gene Annotation and KEGG Pathway Analysis

Differentially methylated CpG sites were annotated to an imported prairie vole genome using a Homer suite tool [56]. Differential CpGs located in gene promoters and gene bodies were assigned to the corresponding genes. Gene promoters were defined as the region from 2000 base pairs upstream to the transcription starting site (TSS) of each gene. Gene bodies referred to the region from TSS to 1000bp downstream of transcription termination site (TTS). The associated annotated gene names, were evaluated for over-represented KEGG pathways, as mentioned above, using the orthologous mouse gene annotation, GRCm38 [41] from Ensembl’s Biomart database.

#### RNAseq and RRBS Correlation Analysis

Differentially expressed genes with differentially methylated CpG sites were chosen for RNAseq and RRBS correlation analysis. The analysis only included the differentially methylated sites located at gene promoters and gene bodies, and their correlation with transcription changes were done separately. First, for all differentially expressed genes that have DNA methylation changes within the gene body, they were separated into eight quantiles (i.e., 12.5% consecutive increments) according to the value of log_2_ fold gene expression change. Furthermore, for all differentially expressed genes that have differentially methylated sites in their promoters, they were divided into four quartiles (i.e., 25% consecutive increments) based on the value of log2 differential gene expression fold changes. Finally, for either gene body or gene promoter analysis, spearman’s correlation was applied to examine any significant correlation between transcription and DNA methylation changes within each of the 8 quantiles or 4 quartiles, respectively.

Furthermore, we evaluated the over-represented biological pathways on all genes that had both transcription change and differential DNA methylation in promoter or gene body regions. The analysis was done in each of the four comparisons separately (i.e., “Attacker” vs “Parental”, “Attacker” vs “Father”, “Parental” vs “Father”, and “Father” vs “Mother”), but without segmenting the differential expression data into quantiles. They were evaluated for over-represented KEGG pathways using the annotated mouse gene IDs as mentioned above. These pathways were imported into Cytoscape and were used to construct a similarity network within the EnrichmentMap [57] plugin. The ClusterMaker [58] plugin was used to cluster the network using the affinity propagation algorithm [59] by identifying “exemplars” or highly connected nodes. The resulting clustered network was visualized using Cytoscape v3.9.1 [53]. Cluster labels were assigned using the AutoAnnotate [60] plugin which takes network node information and automatically assigns cluster labels according to a word tag cloud.

## RESULTS

### Behavior

During the pup exposure test, both “Mother” and “Father” groups displayed similarly high levels of parental behaviors. Some virgin males showed spontaneous parental behaviors including licking and grooming the conspecific pup, huddling over, pup carrying, as well as nest building. These voles were assigned to the “Parental” group (Fig. 1A). In contrast, the other virgin males that demonstrated aggression toward the conspecific pup as indicated by a short latency to attack, were assigned to the “Attacker” group (Fig. 1B). Although “Parental” virgin male voles were similar as “Mother” and “Father” in overall parental behavior frequency and duration, they appeared to be different in pup huddling frequency (one-way ANOVA, F(2,18) = 6.812, p = 0.006, Tukey’s HSD p = 0.005, Fig. 1C, Table S1) and duration (F(2,18) = 4.453, p = 0.027, Tukey’s HSD p = 0.023, Fig. 1D, Table S1). Furthermore, we found the non-parental behaviors were generally comparable among groups (Figs. 1E, 1F), except “Parental” virgin male voles demonstrated a preference for resting, a self-serving behavior (rest frequency, “Parental” vs “Mother”, F(2,18) = 6.191, p = 0.009, Tukey’s HSD p = 0.008, Table S1). We also tested group effects on rest duration and olfaction duration, which were significant for the overall model, but had no statistical significance under Tukey HSD post-hoc conditions (Table S1).

**Fig. 1:**
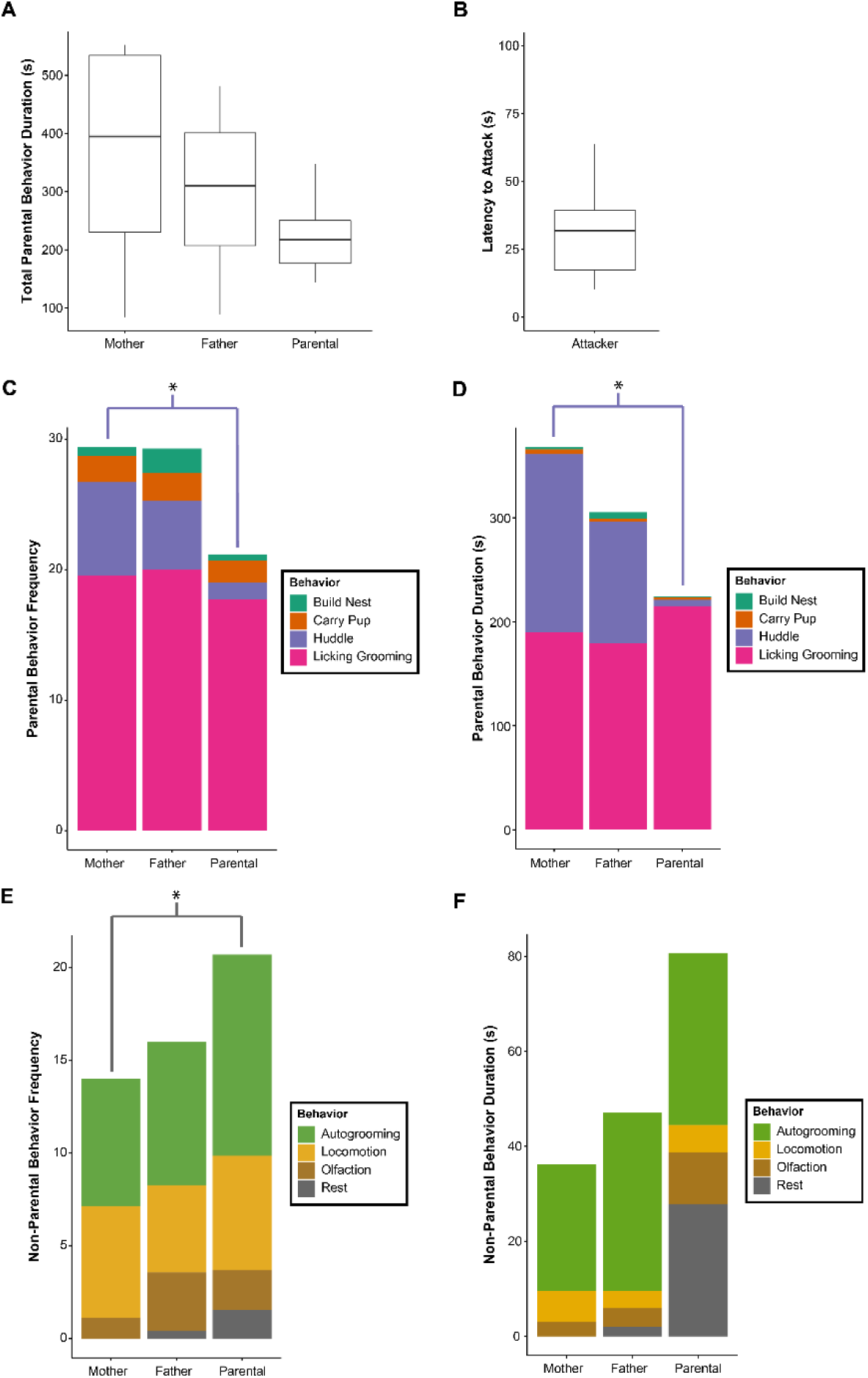
Paternal behavioral dichotomy in virgin male prairie voles. **A**. Summary of the duration of total parental behavior in seconds for each of the three experimental groups (“Mother”, “Father”, “Parental”). **B.** Latency to attack in seconds by the “Attacker” virgin males. In **A** and **B**, whiskers of boxplot represent the ranges of each group, the upper and lower bounds of the boxes represent inter-quartile range (IQR) from quartile 1 and quartile 3, and the horizontal line represents the median. **C,D,E,F**. Behavioral measurements of build nest, carry pup, huddle, licking grooming (**C**, **D**), and autogrooming, locomotion, olfaction, rest (**E**, **F**), in frequency (**C**, **E**) and duration (**D**, **F**) of “Mother”, “Father”, and “Parental” groups. Stacked barplots are color coded according to the legend. Data was analyzed with a one-way ANOVA with TukeyHSD post-hoc correction. *represents p-value < 0.05.

### Transcriptome

We found numerous differentially expressed genes (DEGs) in the four pair-wise comparisons of “Attacker” vs “Parental”, “Attacker” vs “Father”, “Parental” vs “Father”, and “Father” vs “Mother” (Figs. 2A, 2B, Table S2). Though there are relatively fewer gene expression changes when comparing across pair-bonding experience or biological sex (N=734 in “Parental” vs “Father”, N=585 in “Father” vs “Mother”, Fig. 2B, Table S2), we found higher numbers of differentially expressed genes when “Attacker” group is compared to “Parental” or “Father” group (N=1,313 and 1,063, respectively, Fig. 2B, Table S2), which indicates a more deviated DG transcriptome in “Attacker” virgin males, compared to the other groups of voles. Furthermore, both of these comparisons yielded more up-regulated genes (“Attacker” vs “Parental”: N=518, “Attacker” vs “Father”: N=305. Table S2) than down-regulated ones (“Attacker” vs “Parental”: N=35, “Attacker” vs “Father”: N=47; Table S2). We then investigated the over-represented KEGG pathways to obtain biological insights of the transcriptome change. We found the largest number of over-represented pathways (N=22) in the “Attacker” vs “Parental” comparison, and a moderate number of enriched pathways in the “Attacker” vs “Father” or “Parental” vs “Father” comparison (N=10, or 4, respectively) (Table S3). In contrast, we identified no over-represented pathway in the “Father” vs “Mother” comparison. Though generally more enriched in the “Attacker” vs “Parental” comparison, a number of these pathways are shared between the “Attacker” vs “Parental”, and “Parental” vs “Father” comparisons, such as “ECM-receptor interaction” and “protein digestion and absorption” (Figs. 2C, 2D, and Table S3). This suggests their expression changes may be implicated in parental behavioral variations in males. Furthermore, a set of genes that includes *Fzd*, *Tcf* and *Wnt* members were consistently detected in the enriched pathways specific to the “Attacker” vs “Parental” and “Attacker” vs “Father” comparisons, but not the other comparisons (Table S3). Though these genes are primarily Wnt signaling molecules, they may also be associated with other KEGG enriched pathways (e.g., Hippo signaling, cancer related pathways. Figs. 2C, 2D, and Table S3). Furthermore, we found the overlapping pathways between the “Attacker” vs “Parental” and “Attacker” vs “Father” comparisons do not include the exact same DEGs. There are 32 overlapping DEGs shared within the overlapping KEGG pathways between the two analyses (Fig. 2E). Among them, many are selectively enriched in a few pathways that include Wnt signaling. For example, *Tcf7* and *Tcf7L2*, transcription factors modulating canonical Wnt signaling pathway output, are both differentially expressed within the “Attacker” vs “Parental” (*Tcf7L2*: log_2_FC = 1.06, p-value = 6.81×10-5; *Tcf7*: log_2_FC = 0.77, p-value = 0.0146, Fig. 2F, Table S2) and “Attacker” vs “Father” comparisons (*Tcf7L2*: log_2_FC = 0.635, p-value = 0.0239; *Tcf7*: log_2_FC = 0.712, p-value = 0.037, Fig. 2F, Table S2). We also found the main target for the canonical Wnt and BMP signaling *Lef1* is up-regulated in both comparisons (*Lef1*: log_2_FC = 0.563, p-value = 0.011, “Attacker” vs “Parental”; *Lef1*: log_2_FC = 0.489, p-value = 0.037, “Attacker” vs “Father”, Fig. 2F, Table S2). Together, these results point to the contrasting transcription signatures associated with the paternal behavioral dichotomy seen in virgin male voles and suggests the involvement of selective biological pathways.

**Fig. 2:**
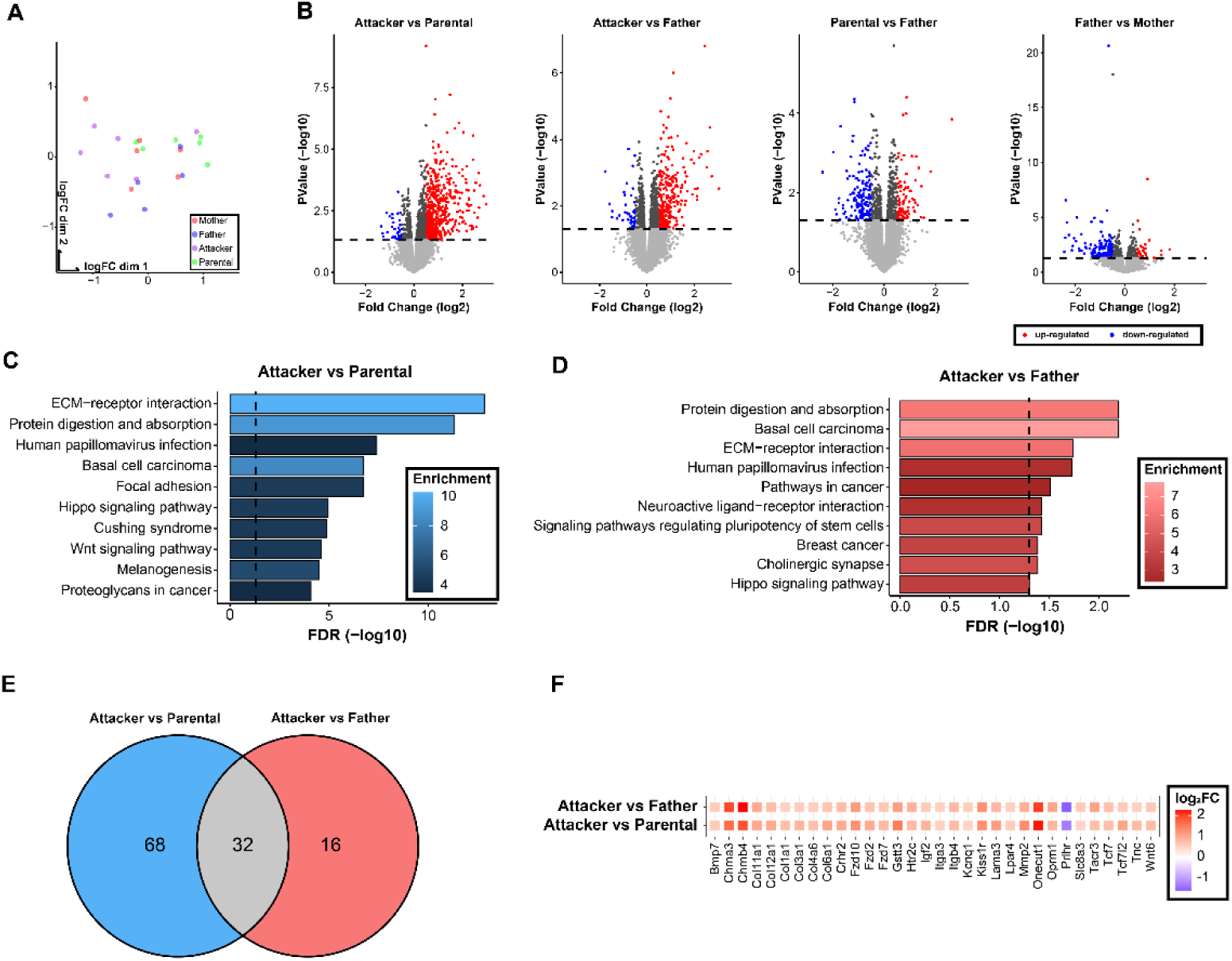
Hippocampal dentate gyrus transcriptome analysis. **A.** Principal component analysis plot showing the variability of twenty-three RNA-seq libraries across all groups (6 Attacker, 6 Parental, 5 Father, 6 Mother). Points are color coded according to experimental group, and labels are dodged to increase clarity. **B.** Volcano plot visualizing differentially expressed genes, in log_2_ transformed Fold Change (log_2_FC) on the x-axis, with −log_10_ adjusted p-values on the y-axis. Up-regulated and down-regulated genes are represented in red and blue, respectively, in the comparisons of “Attacker” vs “Parental”, “Attacker” vs “Father”, “Parental” vs “Father”, and “Father” vs “Mother”, as shown from the left to the right. The dashed line represents –log_10_(0.05). **C, D.** Top ten significantly enriched KEGG pathways resulting from the “Attacker” vs “Paternal” comparison (**C**) and the “Attacker” vs “Father” comparison (**D**). The color gradient for each bar represents the enrichment ratio (observed/expected) with brighter color showing higher enrichment. The x-axis represents the −log_10_ transformed FDR value. **E.** Venn diagram showing the numbers of differentially expressed genes in the enriched pathways of the KEGG pathway over-representation test of “Attacker” vs “Paternal” comparison (blue) and “Attacker” vs “Father” comparison (red), with the grey showing the union of the 32 overlapping genes between the two datasets. These overlapping genes are presented in a dotplot in **F.** with color code assigned to represent the log_2_FC from the RNAseq comparison.

To further characterize the “Attacker” transcriptome, we performed RRHO [47, 48], an unfiltered transcriptome analysis, and found a vast degree of concordant signals between the “Attacker” vs “Parental” and “Attacker” vs “Father” transcriptome comparisons (total 3,723 genes up-regulated, 4,392 genes down-regulated, unfiltered analysis, Fig. 3A and Table S4). However, virtually no discordant signal was detected in the same analysis (only 1 gene that is down in “Attacker” vs “Parental” comparison is up in “Attacker” vs “Father” comparison; and no overlap between up-regulated genes in “Attacker” vs “Parental” comparison and down-regulated genes in “Attacker” vs “Father” comparison; Fig. 3A and Table S4). This further supports the notion of a profoundly contrasting transcriptome in “Attacker” virgin males when compared to “Parental” and “Father” groups, which shared broad similarities.

**Fig. 3:**
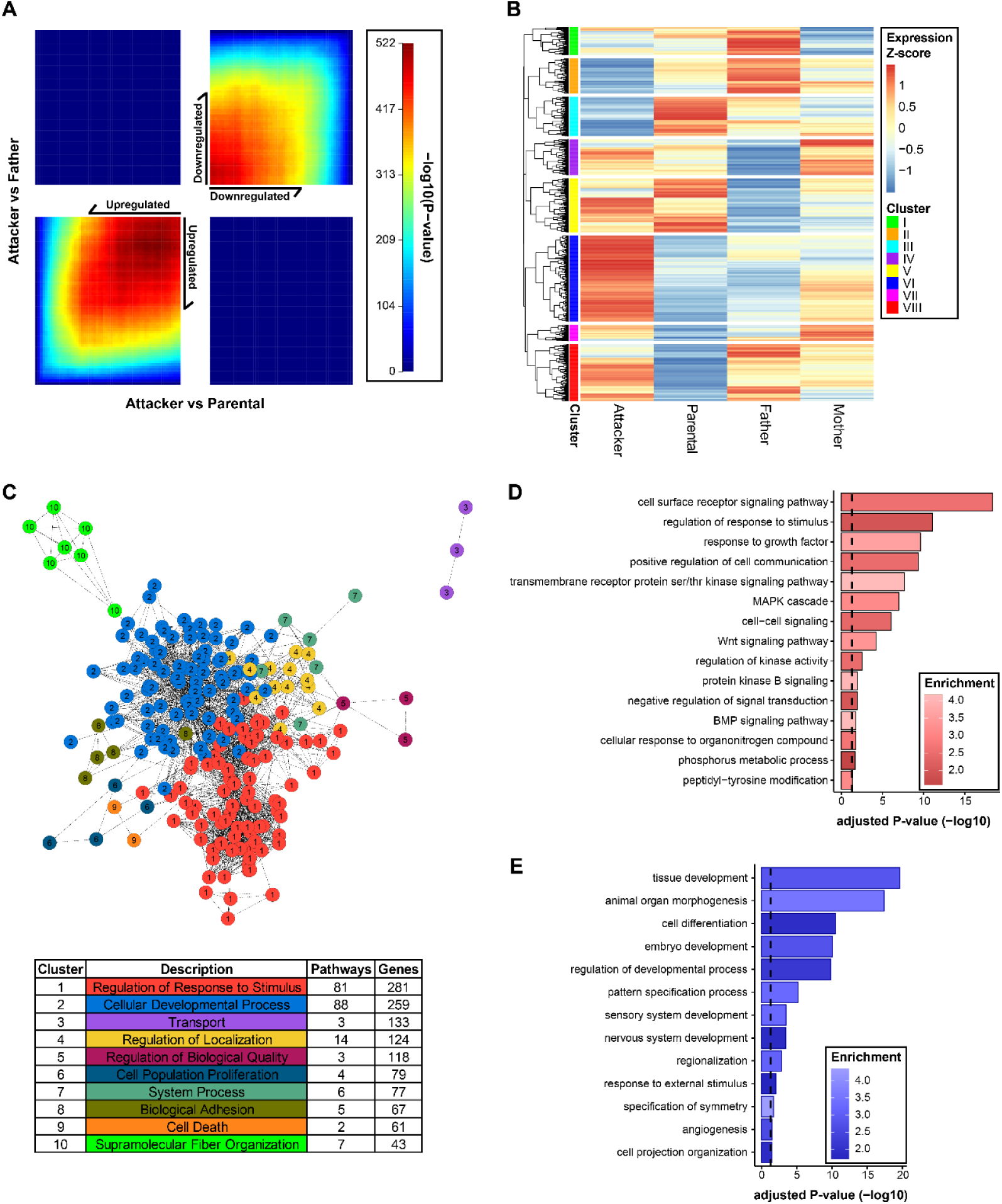
A more deviated transcriptome in Attacker virgin males. **A.** RRHO analysis of all expressed genes in the “Attacker” vs “Parental” and “Attacker” vs “Father” comparisons, demonstrating either a discordant (top left and bottom right quadrants), or a concordant relationship (top right and bottom left quadrants). The overlap is made in a whole-transcriptome and threshold-free manner, where each pixel represents an overlap of the ranked lists. The color of each pixel represents the Benjamini-Yekutieli adjusted −log_10_(p-value) of a hypergeometric test, with warmer colors reflecting more significance. **B.** Clustered heatmap of all differential genes in each of the four comparisons. The analysis was done by the “pheatmap” software, with rows representing z-scores of normalized gene expression among the four groups in the analysis. The eight clusters (I-VIII) were generated using k-means clustering based on the normalized z-score of genes. **C.** GOMCL cluster analysis of genes in heatmap cluster VI (**B**). Significantly over-represented biological process gene ontologies were represented in 10 clusters with reduced redundancy. The chart describes the ten simplified clustered pathways with the respective number of pathways included and the number of unique genes within each pathway to the right. Representative GO Biological Process Pathways of GOMCL cluster 1 and cluster 2 are displayed in **D** and **E**, respectively. The over-represented pathways were modeled using the hypergeometric test with the whole annotated transcriptome as background. The color gradient of each bar represents the enrichment ratio (observed/expected), with brighter color showing higher enrichment. The x-axis represents −log_10_ transformed FDR values.

To obtain a genomic scale overview of transcriptome changes across all groups, we carried out a nuanced approach by constructing a “Differential Expression Clustering” heatmap analysis that includes all DEGs from any of the four comparisons (i.e., “Attacker” vs “Paternal”, “Attacker” vs “Father”, “Paternal” vs “Father”, “Father” vs “Mother”; Table S2) through plotting their inter-group gene expression z-scores (Table S5). Among the eight clusters (I to VIII) classified through k-means clustering, cluster VI has the highest number of DEGs. They were up-regulated in “Attacker”, which is more discordant from the other three groups that were generally down-regulated (Figure 3B). As cluster VI genes were over-represented in numerous biological process pathways (N=240 over-represented pathways, Table S6), we further performed a “Gene Ontology Clustering” analysis to construct a clustered similarity network with the majority of these over-represented pathways included (N=213) to facilitate our interpretation (Fig. 3C, Table S7). Of the ten clusters (named in Arabic numbers 1 to 10) we derived, clusters 1 and 2 contained the largest set of biological process pathways and the highest number of genes (Cluster 1 = 81 pathways, 281 genes; Cluster 2 = 88 pathways, 259 genes, Fig. 3C, Table S7). Together, clusters 1 and 2 account for close to 80% of biological pathways and about 43% of DEGs in differential expression cluster VI. Within cluster 1 (Fig. 3D, Table S7) that is represented by the parent GO term “regulation of response to stimulus”, we found an enrichment of several signaling pathways, with some also recognized within the aforementioned KEGG pathway analysis (Fig. 2), such as Wnt signaling (Enrichment = 3.42, adjusted p-value = 0.00128, Fig. 3D and Table S7). For cluster 2 analysis that is represented by the parent GO term “cellular developmental process”, a number of development pathways are enriched, such as nervous system development (Enrichment = 1.851, adjusted p-value = 0.000349, Table S7), both positive and negative regulation of cell-differentiation (positive regulation: Enrichment = 2.259, adjusted p-value = 0.0123; negative regulation: Enrichment = 2.85, adjusted p-value = 4.55×10^−5^, Table S7). This may reflect DG’s role in adult neurogenesis [61] that contributes to the paternal behavioral dichotomy.

Beyond cluster VI, in the “Differential Expression Clustering” heatmap (Fig. 3B), clusters II and III also displayed an opposing transcription pattern in “Attacker” compared to the other three groups, with genes in “Attacker” group being down-regulated. From them, numerous gene ontology terms were over-represented, which included neurogenesis (Enrichment = 2.67, adjusted p-value = 9.02×10^−5^), and synaptic signaling (Enrichment = 5.65, adjusted p-value = 7.48×10^−10^, Table S6), two of the top enriched ontology terms. Together, our results demonstrate a unique biological signaling signature of the DG transcriptome associated with the paternal behavioral dichotomy.

### DNA Methylome

To obtain a molecular insight of parental behavior beyond the transcriptome, we examined DNA methylation in the DG, which may mediate gene expression. By using RRBS methylome profiling [36], we found numerous differentially methylated CpG sites in each of the four pair-wise comparisons with consistently more hypermethylation sites than hypomethylation sites (Fig. 4A, Table S8). Unlike what we found in the transcriptome analysis that “Attacker” vs “Parental” and “Attacker” vs “Father” comparisons had the most transcription changes than the other comparisons, the highest number of differentially methylated CpG sites was found in “Parental” vs “Father” and “Father” vs “Mother” comparisons (total number in 34,453, and 34,468, respectively). In contrast, “Attacker” vs “Parental” and “Attacker” vs “Father” comparisons have substantially lower number of methylation changes (24,212 and 28,549 differentially methylated CpG sites, respectively. Fig. 4A, Table S8). This may be explained by the plausible involvement of DNA methylation in sex difference or sexual experience. In addition, we found that most of the differentially methylated CpG sites are located in intergenic and intronic genomic regions (around 50% and 30%, respectively, in each comparison; Fig. 4B, Table S8), and around 3% of differentially methylated CpG sites reside are at promoters (Fig. 4B).

**Fig. 4:**
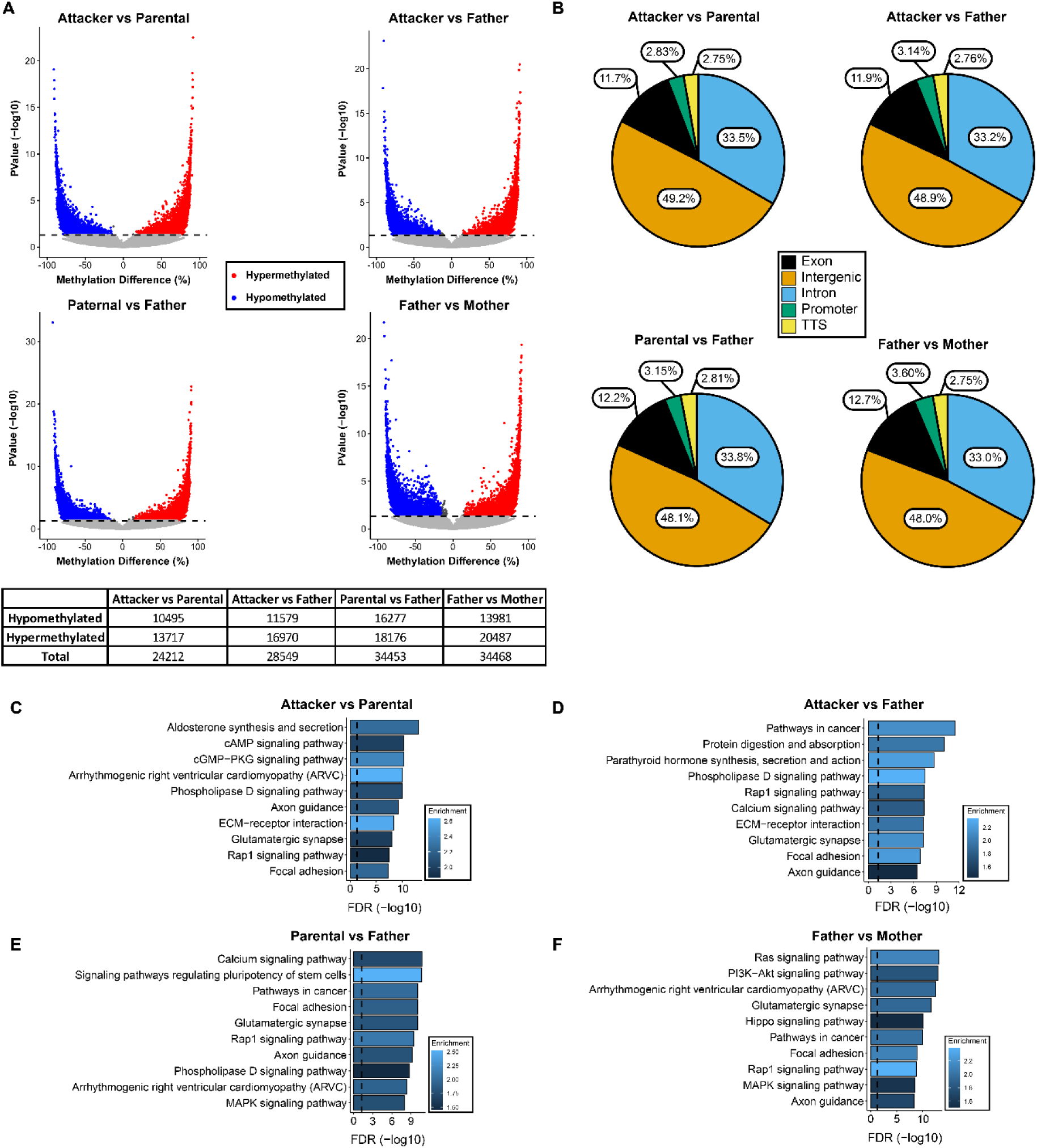
Differential DNA methylation analysis. **A.** Volcano plot visualizing differentially methylated CpG sites. Percentage of methylation difference is represented on the x-axis, while the −log_10_(p-value) is represented on the y-axis. Hypermethylated CpGs and hypomethylated CpGs are represented in red and blue, respectively. The horizontal dashed line corresponds to −log_10_(0.05) to represent a threshold for significance. The chart on the bottom demonstrates the numbers of differentially methylated sites in each comparison. **B.** Pie charts represent the genomic feature distribution of the differentially methylated CpG sites from each comparison. Labels within each section represent the % of differentially methylated CpG sites that fall within these features. TTS stands for transcription termination site. **C, D, E, F.** The top 10 ranked over-represented KEGG pathways of genes that contain differentially methylated CpG sites within their boundary (from 2,000 bp upstream of the transcription start site to 1000 bp downstream of the transcription termination site) for the comparisons of “Attacker” vs “Parental” (**C**), “Attacker” vs “Father” (**D**), “Parental” vs “Father” (**E**), and “Father” vs “Mother” (**F**). In each of the panels, the vertical dashed line represents an FDR cutoff of 0.05, and the color represents the enrichment ratio of the pathways.

To explore the functional implication of this vast array of DNA methylation changes, we examined the pathways over-represented by genes containing differentially methylated CpG sites in promoter or gene body regions (Table S9). We found that over 100 KEGG pathway terms are enriched in each of the four comparisons and many of them are overlapping. By examining the top 10 enriched pathways (Figs. 4C-4F), none appears to be unique to each comparison. Instead, over 80 pathways are over-represented in all four comparisons (Fig. S1A, Table S9). These include “ECM-receptor interaction” and “protein digestion and absorption”, the top two common pathways identified in the aforementioned transcriptome comparisons (Figs. 2C, 2D, and Table S3), which suggests a canonical transcription regulatory role of DNA methylation in paternal behavior. “Oxytocin signaling”, a key pair-bonding pathway that is known to subject to DNA methylation modulations [23], is also enriched in all comparisons (Fig. S1A). Additionally, “Rap1 signaling”, “glutamatergic synapse”, “axon guidance”, “PI3K-Akt signaling”, “calcium signaling”, “cAMP signaling”, “cholinergic synapse”, “Ras signaling”, “GnRH signaling”, “thyroid hormone synthesis” are all notable overlapping pathways, to name a few (Fig. S1A). Furthermore, we found 60 pathways over-represented in one, two, or three, but not four, comparisons (Fig. S1B, Table S9), with over half of them not enriched in the “Attacker” vs “Parental” comparison, such as “neurotrophin signaling pathway” and “VEGF signaling pathway”. Among them, a dozen pathways are only enriched in one comparison (e.g., P53 signaling in “Attacker” vs “Father”, vasopressin in “Father” vs “Mother”), but their enrichments are not the highest in accordance to FDR values (Fig. S1B). In the meantime, we found some immune related pathways appearing to be enriched in “Father” related comparisons, which include “NF-kappa B signaling”, “T cell receptor signaling”, “B cell receptor signaling”, “Fc signaling”, to name a few (Fig. S1B).

### Transcriptome-Methylome Correlation

The numerous differential methylation pathways that are mostly overlapping across comparisons may suggest DNA methylation’s canonical roles widely associated with parental behaviors. Therefore, the DNA methylation changes may represent a spectrum of molecular signatures of the parental behavior variations as indicated by the inter-group behavioral examinations (Fig. 1). Given the regulatory role of DNA methylation in gene transcription, we then moved forward to explore any association between DNA methylome and transcriptome alterations.

Although in each of the four pair-wise comparisons, we did not find an overall correlation between DNA methylation and transcription for genes that had both transcription and DNA methylation changes, we noticed a significant correlation within subsets of genes when they were separated into quantiles based upon transcription levels. The DEG quantile classification was done in each pair-wise comparison individually. They were further separated for DNA methylation changes in promoters or gene bodies, respectively, to potentially differentiate DNA methylation’s effect in these two genomic regions. More specifically, in each pair-wise comparison, all differentially expressed genes that have differentially methylated sites in their promoters were divided into four quartiles (i.e., 25% consecutive increments) based on the value of log_2_ differential expression change. In addition, all differential expressed genes that have DNA methylation changes within the gene body were separated into 8 quantiles (i.e., 12.5% consecutive increments) according to the value of log_2_ fold gene expression change (Figs. 5A-H). We then examined each of these quantiles individually for a correlation between transcription and DNA methylation changes (Tables S10, S11). We found a few correlations, with some in the quantiles with extreme expression changes. For example, a positive correlation was found between differential expression and gene body DNA methylation in highest expressed quantile 8 of “Parental” vs “Father” comparison (cor = 0.253, p-value = 0.011, Fig. 5I bottom-left, Table S10), and quantile 2 of “Father” vs “Mother” comparison (cor = 0.211, p-value = 0.027, Fig. 5I bottom-right, Table S10). In addition, there were a negative correlation between transcription and gene body DNA methylation in the “Attacker” vs “Parental” quantile 8 comparison (cor = −0.187, p-value = 0.011, Fig. 5I, top-left, Table S10), and a positive correlation between differential transcription and promoter DNA methylation changes in one of the highest expressed quartiles in the “Attacker” vs “Father” comparison (quartile 3: cor = 0.683, p-value = 0.014, Fig. 5J, top-right, Table S10). Given the relative uniformity of cellular composition in DG, the enrichments of DNA methylation changes at the quantiles of genes with extreme expression changes may indicate DNA methylation as a canonical transcription regulatory mechanism existing in the majority of cells in DG to mediate paternal behavior.

**Fig. 5:**
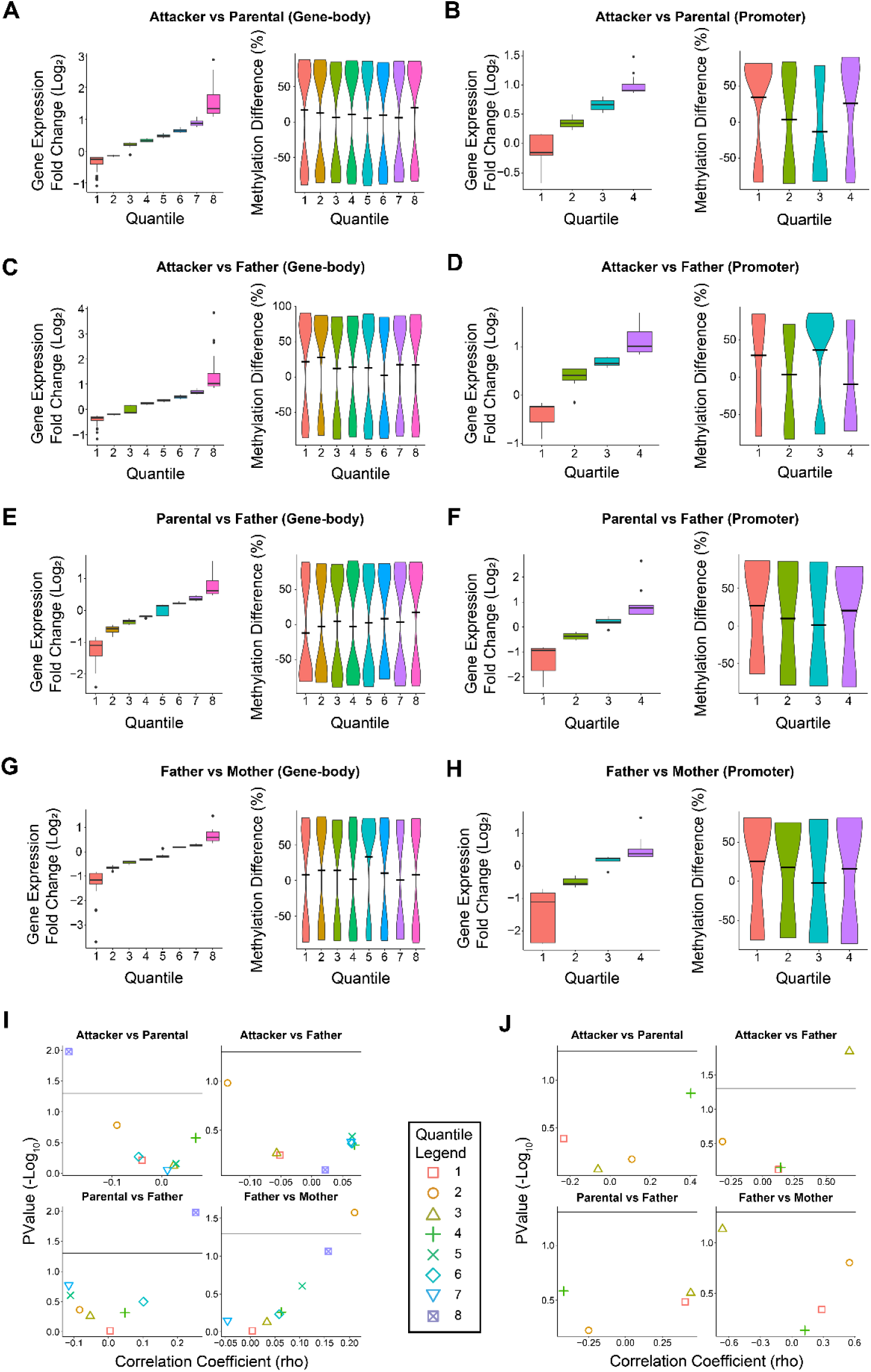
Correlations between gene expression and DNA methylation changes. **A.** In “Attacker” vs “Parental” comparison, genes that have both differential expression and differentially methylated CpG sites within their gene boundary were divided into eight quantiles based upon the value of log_2_ fold gene expression change. The left plot represents the log_2_ fold change of the differentially expressed genes, with the horizontal line represents the median of each group. The right plot indicates DNA methylation differences of each of the eight quantiles. The width of the violin plot indicates the proportion of datapoints at y-axis values. The black line represents the mean value of the methylation differences. **B.** In “Attacker” vs “Parental” comparison, the analysis is done in the same manner as in **A** for differentially expressed genes that have DNA methylation changes in their promoters, except all genes were separated into 4 quartiles. For pairs **C** & **D, E** & **F**, and **G** & **H**: they represent the same type of analysis as in **A** & **B**, but for “Attacker vs Father”, “Parental vs Father”, and “Father vs Mother” comparisons respectively. **I** & **J.** A scatter plot of spearman correlation coefficients and the associated p-values from the correlation analysis of quantiles in the four comparisons of the analysis with their labels listed above each portion of the chart. The horizontal line represents −log_10_(0.05) as a threshold of significance. Panels in **I** correspond to the eight quantiles in **A**,**C**,**E**, and **G,** while panels in **J** correspond to the four quartiles in **B**,**D**,**F**,**H**.

Within the significantly correlated quantiles, we recognized a number of functional meaningful genes, which included the ones belonging to the Wnt signaling pathway. For example, *Dkk3*, a secreted inhibitor of Wnt signaling, was slightly under-expressed while having a hypermethylated CpG in the promoter region of “Attacker” (log_2_FC = −0.264, differential methylation = 0.743, “Attacker” vs “Parental”, Table S11), which is consistent with the repressive role of DNA methylation on gene transcription at gene promoters. Furthermore, *Fzd10*, a membrane bound Wnt receptor, was up-regulated with a hypomethylated CpG within intron 1 (log_2_FC = 1.280, differential methylation = −0.613, “Attacker” vs “Parental”, Table S11). In addition, there were two gene body hypermethylated sites in the upregulated gene *Enpp2* (log_2_FC = 1.862, “Attacker” vs “Parental”, Table S11), whose protein product can function as a phosphodiesterase to catalyze the production of lysophosphatidic acid (LPA), which is known to activate the β-catenin pathway[62, 63]. Together these results implicate that DNA methylation selectively impacts certain regulatory systems to affect paternal behaviors.

To obtain a systemic biological interpretation of the genes that have both transcription and DNA methylation changes, we analyzed the KEGG pathways over-represented by these genes (Fig. 6, Table S12). We found the highest number of enrichment terms in the “Attacker” vs “Parental” comparison (N=65), followed by the “Attacker” vs “Father” comparison (N=16, Figs. 6A, B, Table S12). In contrast, in “Parental” vs “Father” and “Father” vs “Mother” comparisons, only two and three pathways were enriched, respectively, and none of them was unique to any comparison (Fig. 6C, Table S12). These suggest that DNA methylation may play a main regulatory role on gene expression associated with the aggressive paternal behavior. Moreover, among the 65 enrichment terms in the “Attacker” vs “Parental” comparison, 49 pathways were not over-represented in other comparisons (Fig. 6A, Table S12), which may indicate their unique contributions to paternal behavioral dichotomy in virgin male voles. Using affinity propagation clustering, implemented in the Cytoscape EnrichmentMap plugin, [57–59] further illustrates that the majority of these pathways (N=40) are highly connected under one single main functional cluster “Wnt signaling pathway”, which also has a substantial overlap with “Thyroid hormone synthesis” (Fig. 6A, Table S12). Although the “Father” group voles have a more similar parental behavioral phenotype to the “Parental” group (Fig. 1), who also share an overall more similar transcriptome when compared to the “Attacker” virgin voles (Figs. 2, 3), the KEGG pathway analysis on genes with both transcription and DNA methylation changes in the “Attacker” vs “Father” comparison did not lead to any many unique pathways as in the “Attacker” vs “Parental” comparison. Instead, only two pathways were uniquely enriched in the “Attacker” vs “Father” comparison, with both of them addiction related (“cocaine addiction” and “amphetamine addiction”, Fig. 6B, Table S12). In the meantime, we found the majority of the pathways shared in two or more comparisons (11 out of 16, Fig. 6C) were only over-represented in the “Attacker” vs “Parental” and “Attacker” vs “Father” comparisons.

**Fig. 6:**
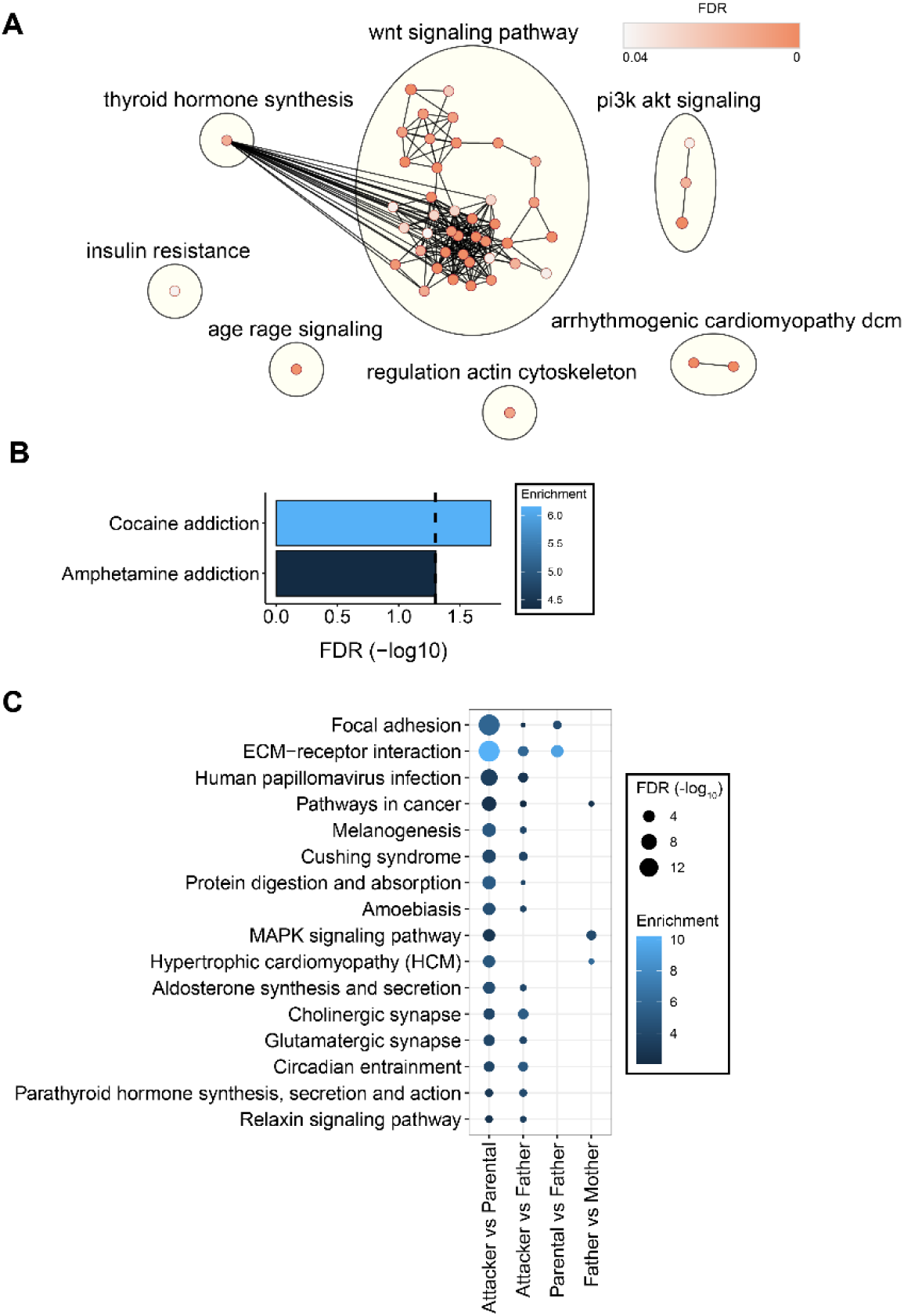
Pathway analysis of genes with both transcription and methylation changes. **A.** The unique over-represented KEGG pathways in the “Attacker” vs “Parental” comparison. KEGG Pathway analysis was done after clustering using an affinity propagation algorithm implemented in the Cytoscape tool. Seven main functional clusters (e.g., Wnt signaling pathway) were identified (yellow circles), each containing one or more connected KEGG terms (nodes). The color of each node represents the FDR value from the over-representation analysis, while the edges of the network represent similarity between the clustered KEGG pathways. **B.** The unique over-represented KEGG pathways in the “Attacker” vs “Father” comparison. The vertical dashed line represents FDR of 0.05, while the color represents the enrichment ratio of the over-representation test. **C.** The shared over-represented KEGG pathways from the comparisons listed on the x-axis. Size of each dot represents the −log_10_ transformed FDR values, while the color represents the enrichment ratio from the over-representation test. The missing datapoints show that the respective pathway was not found to be over-represented in the comparison listed.

## DISCUSSION

In this study, we examined the dentate gyrus (DG) transcriptome and methylome to obtain molecular insights of parental behaviors of prairie voles following a pup-exposure paradigm. In this behavioral model, both father and mother voles display biparental care, whereas virgin male voles demonstrate parental behavioral dichotomy with some behave parentally and the rest show aggression. Though the prairie vole genome is still under construction with large portions of chromosomes isolated on independent genome scaffolds which poses a challenge for genomic research, our exploratory study has led to some interesting findings that support a DNA epigenetics based molecular underpinning of paternal behavioral dichotomy.

We found that spontaneously aggressive virgin male prairie voles (“Attacker”) exhibited unique molecular signatures (both transcriptome and methylome changes) compared to spontaneously parental virgin male voles (“Paternal”). Though variable gene transcription patterns have been reported in other brain regions[64], we found profound genome-wide gene expression changes in the DG of “Attacker” virgin males, when compared to either “Parental” or “Father” voles, which were more comparable. The differential genes from these analyses are selectively enriched in a number of biological pathways that suggest their functional implications in parental behavior. Notably, we found expression changes of several Wnt signaling molecules that are associated with the parental behavior variance. Wnt signaling is a highly conserved pathway that plays fundamental roles in development and homeostasis [65–68]. Accumulating evidence supports its major roles in neural development, synaptic plasticity, as well as brain diseases [69]. Wnt signaling generally includes the canonical and noncanonical pathways. The canonical pathway is usually referred as the “Wnt/β-catenin pathway” due to its dependence on the stabilization of β-catenin. The binding of Wnt ligands to the cell surface Frizzled (Fzd) receptors and LRP5/6 co-receptors will endocytose this complex and inhibits the β-catenin destruction complex that consists of GSK-3β. This leads to increased levels of β-catenin and its translocation to the nucleus to mediate gene expression through the interaction with TCF/LEF transcription factors. In addition, the silencing of the canonical pathway may be carried out through the activation of the β-catenin destruction complex by Wnt/β-catenin inhibitor DKK. In contrast, the non-canonical Wnt signaling pathway functions without β-catenin or GSK-3β as an intermediate molecule. It is triggered by Wnt ligand binding to the Fzd receptor and coreceptors, which then activates either the Wnt/PCP (planar cell polarity) (also named Wnt/JNK) pathway or the Wnt/calcium pathway, two main downstream branches of the non-canonical Wnt signaling. The non-canonical pathways play significant roles in cytoskeleton remodeling, synaptic plasticity, and axon guidance [69]. The identification of various Wnt signaling molecules (e.g., *Wnt*s, *Fzd*s, *Dkk*,*Tcf/Lef*s), and the enrichments of related pathways (e.g., axon guidance, calcium signaling, synaptic signaling, etc.) in our differential transcriptome analyses suggest a broad implication of Wnt signaling in parental behavior modulation that likely includes both canonical and non-canonical pathways. Given the dentate gyrus is a major adult neurogenesis site and Wnt signaling’s involvement in both neurogenesis [70, 71] and mature brain synaptic plasticity, it remains to be addressed at what stage Wnt signaling is involved in parental behavior.

In order to further evaluate the degree of gene expression changes in a whole transcriptome manner, we utilized a modification of a hypergeometric overlaps test RRHO. We found a great deal of similarities between the unfiltered transcriptome comparisons of “Attacker” vs “Parental” and “Attacker” vs “Father”, which provides a novel molecular basis of the paternal behavioral dichotomy. Notably, we found DNA epigenetic modification enzymes, *Dnmt3a* and *Tet1,* have concomitant transcription changes in “Attacker” vs “Parental” and “Attacker” vs “Father” comparisons. DNA methyltransferases (DNMTs) catalyze DNA methylation through the covalent addition of a methyl group to cytosine nucleotide, which can be altered through a series of oxidation reactions mediated by Ten-eleven translocation (TET) methylcytosine dioxygenases, that may ultimately lead to unmethylated cytosines [72–74]. In the future, it will be interesting to identify if any DNA methylation enzyme has expression changes in specified cell types.

Though a few studies have evaluated DNA methylation in prairie voles, they all focused on individual genes, particularly nonapeptides (e.g., vasopressin and oxytocin) [20, 75, 76]. To our knowledge, our study is the first to investigate DNA methylation in prairie vole brains at a genomic scale. We found the differential methylation loci associated genes are enriched in a broader range of KEGG pathways with many not represented in the differential transcriptome analysis. Compared to the transcriptome examination, epigenome profiling may demonstrate molecular changes in a multi-dimensional manner to reflect not only past experience, but also future inducibility. It may also capture alterations in subpopulations of cells that are below the detection threshold of a bulk tissue RNAseq analysis. Likely, some of the DNA methylation changes impact transcriptome outputs in a defined cell population that were unable to be identified in our whole DG tissue RNAseq examination. In addition, the RRBS sequencing we applied in this study only surveys a portion of the DNA methylome [77]. If the RRBS results reflect a representative DNA methylome overview, more widely distributed methylation changes are expected to occur across the whole genome. This will be interesting to explore in the future, which demands significant financial and bioinformatic endeavors. When we performed the methylome pathway analyses, we limited them to differentially methylated sites at gene promoters and gene bodies. The reason to exclude DNA methylation changes at distal intergenic regions, which account for a major subgroup of differentially methylated sites, is because it remains challenging to identify their target genes. Often times, the intergenic regulatory DNA elements bypass their nearby genes to modulate transcription of genes located a long linear distance away through three-dimensional looping [78]. How the higher order genome is organized in prairie vole DG is unknown.

While all above factors may explain the variable findings between our transcriptome and DNA methylome analyses, it is intriguing to see several overlapping pathways that are enriched with both DNA methylation and transcription changes, such as “ECM-receptor interaction” and “protein digestion and absorption”. This suggests DNA methylation may modulate parental behavior associated gene transcription changes in selective pathways. Particularly, we have found that genes with both methylation and transcription changes in the “Attacker” vs “Parental” comparison are profoundly enriched in the “Wnt signaling” cluster (Fig. 6A). Though growing evidence has indicated Wnt signaling’s role in neural development and function in recent years, our integrated transcriptome and DNA methylome examinations suggest a novel function of Wnt signaling in parental behavior, particularly paternal behavioral dichotomy. Furthermore, from the same integrated analysis, we have noticed several other interesting candidate pathways, such as cytoskeleton, cholinergic signaling, immune signaling, that also deserve further investigation. By the high-throughput nature of our genomic approaches, our datasets should serve as a valuable reference for future relevant researches.

In summary, our study has provided an unprecedented integrated view of dentate gyrus transcriptome and DNA methylome in prairie voles, which are associated with the parental behavior differences. The significant correlation between DNA methylation and gene expression in selective biological pathways illustrates a novel role of DNA methylation in parental behavior.

## Data Availability

Next generation sequencing files and processed data for both RNAseq and RRBS datasets is available at the NCBI Geo under the GSE214799 superseries.

## Author contributions

The study was conceived by Z.W., N.J.W., and J.F. Y.L. performed animal experiments. Y.L., G.J.K., N.J.W., performed behavioral data analysis. J.M.C. prepared sequencing libraries. N.J.W. performed bioinformatic sequencing data analysis with feedback from other authors. N.J.W., Z.W., and J.F. prepared the manuscript. All authors reviewed and approved the manuscript.

## Acknowledgements

This work was supported by National Institutes of Health grants (R01MH108527 and R21MH111998 to ZW, DP1DA046587 and R01DA046720 to J.F.). J.M.C was a recipient of FSU Neuroscience Fellowship and FSU Legacy Fellowship.

## List of Supplementary Materials

Supplemental Table 1: Behavioral Analysis

Supplemental Table 2: Differentially Expressed Genes

Supplemental Table 3: Over-represented KEGG Pathways from DEGs

Supplemental Table 4: RRHO Gene List

Supplemental Table 5: Normalized Gene Counts for Clustering

Supplemental Table 6: Cluster-based Gene Ontology Pathways

Supplemental Table 7: GOMCL Ontology Network Analysis

Supplemental Table 8: Differentially Methylated CpG Sites

Supplemental Table 9: Over-represented KEGG Pathways from Differentially Methylated CpG Sites

Supplemental Table 10: Differential Gene Expression and DNA Methylation Quantile Correlation

Supplemental Table 11: Correlated Differential Gene Expression and DNA Methylation Changes

Supplemental Table 12: Overlapping DEGs and Differentially Methylated Sites KEGG pathway enrichment

Supplemental Figure 1: KEGG pathways of differentially methylated CpGs

## Supplemental Figures

**Supplemental Figure 1:**
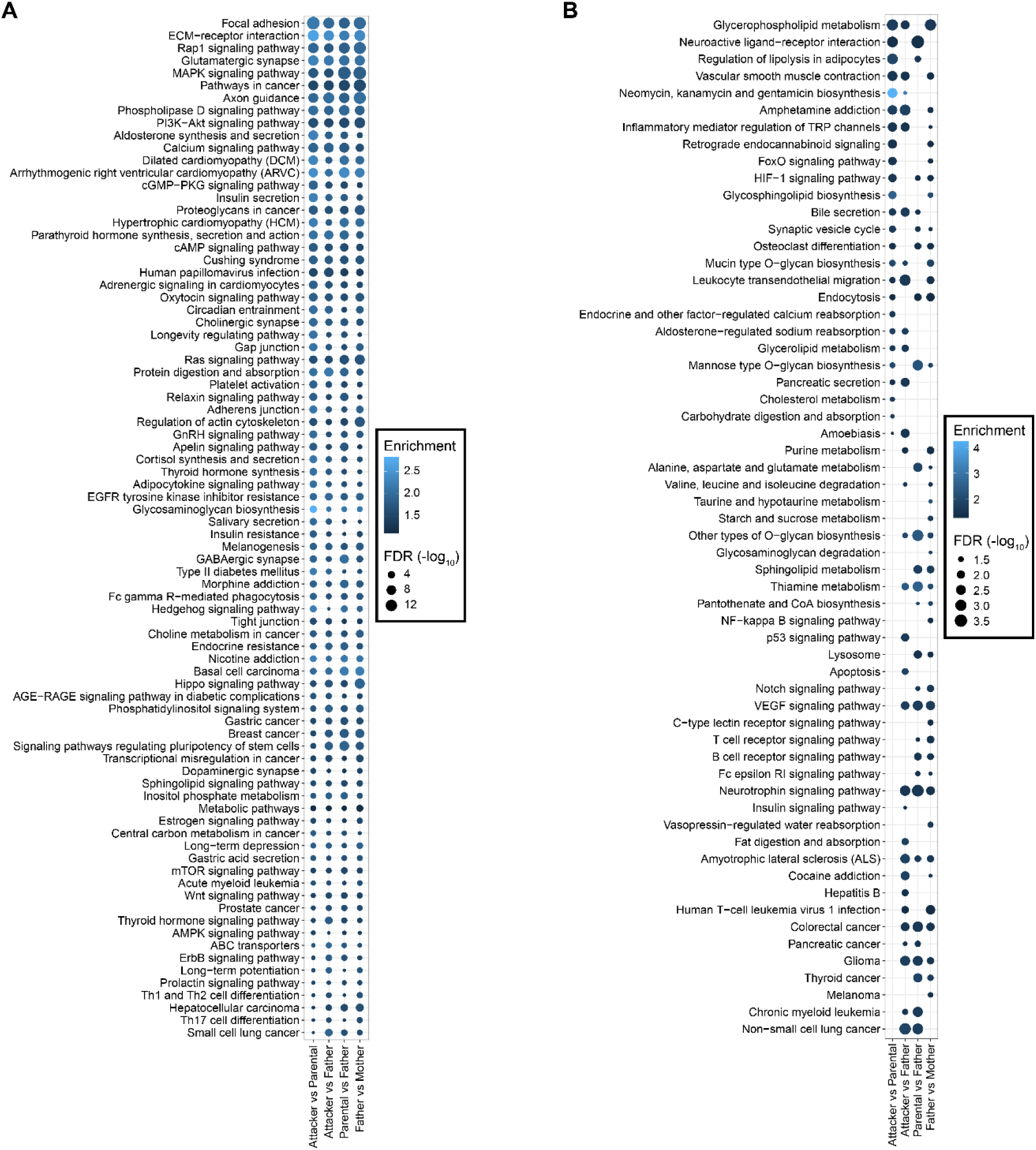
KEGG pathways of differentially methylated CpGs. **A & B.** These dot plots show over-represented KEGG pathways from genes that contained a differentially methylated CpG site within their genic regions from each of the comparisons. This includes (−2000 bp upstream of the TSS to 1000 bp downstream of the TTS). The dot size represents the FDR from the over-representation test, while the color corresponds to the enrichment, or observed / expected values from the test. In **A**, pathways that were shared among all four comparisons are listed, while in **B**, pathways that were enriched in one to three comparisons.

